# Tension-dependent stabilization of E-cadherin limits cell-cell contact expansion

**DOI:** 10.1101/2020.11.20.391284

**Authors:** Jana Slováková, Mateusz Sikora, Silvia Caballero-Mancebo, S.F. Gabriel Krens, Walter A. Kaufmann, Karla Huljev, Carl-Philipp Heisenberg

**Affiliations:** Institute of Science and Technology Austria, Am Campus 1, A-3400 Klosterneuburg, Austria; Institute of Molecular Biotechnology, Dr. Bohr-Gasse 3, 1030 Vienna, Austria; Max Planck Institute of Biophysics, Max-von-Laue-Straße 3, D-60438 Frankfurt am Main, Germany; University of Vienna, Faculty of Physics, Sensengasse 8/9 A-1090 Vienna, Austria

## Abstract

Tension of the actomyosin cell cortex plays a key role in determining cell-cell contact growth and size. The level of cortical tension outside of the cell-cell contact, when pulling at the contact edge, scales with the total size to which a cell-cell contact can grow^1,2^. Here we show in zebrafish primary germ layer progenitor cells that this monotonic relationship only applies to a narrow range of cortical tension increase, and that above a critical threshold, contact size inversely scales with cortical tension. This switch from cortical tension increasing to decreasing progenitor cell-cell contact size is caused by cortical tension promoting E-cadherin anchoring to the actomyosin cytoskeleton, thereby increasing clustering and stability of E-cadherin at the contact. Once tension-mediated E-cadherin stabilization at the contact exceeds a critical threshold level, the rate by which the contact expands in response to pulling forces from the cortex sharply drops, leading to smaller contacts at physiologically relevant timescales of contact formation. Thus, the activity of cortical tension in expanding cell-cell contact size is limited by tension stabilizing E-cadherin-actin complexes at the contact.

## Introduction

For multicellular organisms to form, cells need to establish stable and long-lasting contacts. Consequently, insight into the molecular and cellular mechanisms by which cell-cell contacts are being formed and maintained is central for understanding how multicellularity has emerged in evolution. Adhesion between cells is mediated by various cell-cell adhesion molecules, amongst which Cadherins constitute a key family of adhesion receptors mediating selective Ca^2+^-dependent cell-cell adhesion^3,4^. While much progress has been made in identifying how cadherin adhesion molecules can trigger cell-cell contact formation by binding to each other and associated molecules, such as Catenins^5,6,7^, comparably little is known on how Cadherins transduce forces between cells, and how such force-transduction feeds back on the organization and function of Cadherins at cell-cell contacts.

Generally, Cadherins - and in particular classical Cadherins - are thought to function in cell-cell contact formation in three different ways^8^: (1) they promote cell-cell contact formation by directly lowering interfacial tension at the cell-cell contact zone^9^. How Cadherins achieve this is not yet entirely clear, but the generation of lateral pressure through Cadherin-mediated molecular crowding at the contact zone has been proposed as one potential effector mechanism^10^. (2) Signaling from Cadherins modulate the actomyosin cytoskeleton at the contact site thereby controlling contact growth and maintenance^11^. Effector molecules involved in this process include RhoA and Arp2/3, which both are repressed when Cadherins bind over the contact, and Rac, which is activated upon Cadherin binding^12,13^. (3) Cadherins transduce pulling forces from the contractile actomyosin cortices of the contacting cells over the contact site^1,14,15^. This force transduction allows the contact to grow and reach steady state once those forces are balanced at the contact. Data on cultured cells and primary cells from zebrafish embryos support a critical function of Cadherins in contact expansion. They are thought to disassemble the actomyosin cortex at the contact site and mechanically couple the cortices of the contacting cells at the contact edge^16,1^. These observations led to a model where pulling forces at the contact edge, originating from the contractile cortices of the contacting cells are transduced by Cadherins over the contact and drive contact expansion. Consequently, the level of cortex tension is expected to scale with the size of the contact^1,2^.

Cadherins at cell-cell contacts not only transduce forces between the contacting cells but are also affected by the forces to which they are subjected. Studies on culture cells have provided evidence that tension at Cadherin cell-cell adhesion sites promote Cadherin clustering and reduce their turnover at the contact site^16,17,18^. How tension functions in those processes is not yet fully understood, but tension-induced stabilization of filamentous Actin (F-actin)^19,20,16^ and unfolding of α-Catenin^21^, a key component of the Cadherin adhesion complex^22,23^, are involved. Unfolding of α-Catenin is thought to reveal cryptic binding sites to Vinculin^21,24^, which again enhances binding of α-Catenin to F-actin by simultaneously binding to both molecules^17,25,26,20^.

Yet, how mechanosensing of Cadherin cell-cell adhesion sites affects the function of Cadherins in contact expansion and maintenance remains unclear. To address this question, we have tested how changes in cortex tension affect contact expansion of zebrafish primary germ layer progenitor cells. Contrary to previous expectations^1^, we found that above a critical threshold level of tension, the size of cell-cell contacts becomes smaller rather than bigger. We further found that this restricting influence of cortex tension on contact growth is due to high tension promoting cytoskeletal anchoring of E-cadherin, leading to enhanced clustering and stability of E-cadherin at the contact.

## Results

To test whether the previously reported promoting influence of cortical tension on cell-cell contact expansion^1,2^ represents a generic principle underlying cell-cell contact formation, we sought to analyze its applicability to a wide range of cortical tensions. In order to examine cell-cell contact formation in primary vertebrate cells, we turned to zebrafish germ layer progenitor cells, previously used to study the role of cortical tension in contact expansion^1^. Specifically, we imaged ectoderm progenitor cell doublets either mounted in polymeric wells allowing us to monitor cell-cell contact organization at high resolution (Fig. 1a) or placed on non-adhesive substrates for high-throughput analysis of contact expansion. Consistent with previous observations^1^, we found that reducing cortical tension in cell doublets by exposing them to 10 μM of the Myosin II inhibitor Blebbistatin (Bb) severely reduced expansion of the cell-cell contact surface area (A_c_) (Fig. 1b,c; Fig. S1). Unexpectedly, however, when performing the reverse experiment and strongly increasing cortical tension by exposing cell doublets to 50 nM Lysophosphatidic Acid^27^ (LPA) or over-expressing constitutively active (ca) RhoA in the contacting cells^28^, contact expansion was reduced rather than increased (Fig. 1b,c; Fig. S1). Notably, the effect of LPA on contact expansion became already apparent during the first minute of contact formation (Fig. 1c, Fig. S1), suggesting that high cortical tension restricts contact expansion within seconds after contact initiation. Together, these findings contrast previous observations of cortex tension promoting contact expansion^1,2^.

**Figure 1.**
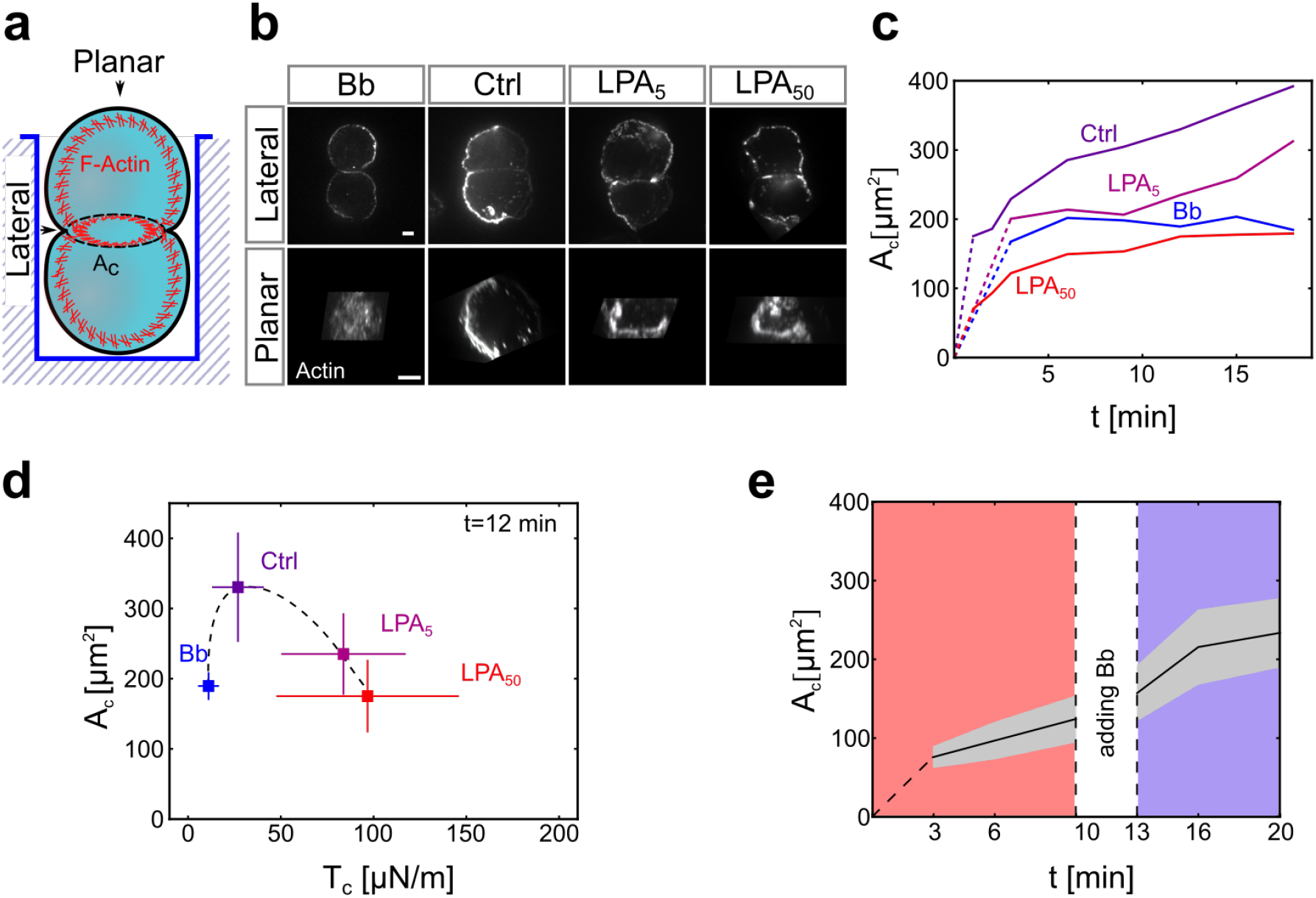
Cortical tension limits contact expansion in cell doublets. (**a**) Schematic of the experimental setup for live imaging of progenitor cell doublets. Doublets were placed in polymeric wells for maintaining their contact within the focal plane while being imaged from the bottom. **(b)** Planar and lateral views of the Actin cell cortex in progenitor cell doublets (10 min contact time) visualised by Phalloidin (F-actin) in control doublets and doublets exposed to Bb (10 μM) or LPA (5, 50 nM). Scale bar, 5 μm. **(c)** Cell-cell contact size (A_c_) as a function of contact time in control doublets, doublets exposed to Bb (10 μM) or different concentrations of LPA (1 - 50 nM). Dotted lines connect contact formation (0 min) with the first time point when data were collected. Error bars (standard deviation) are shown in Fig. S1. (Ctrl) N = 9, n = (1min: 30, 2min: 26, 3min: 30, 6min: 24, 9min: 24, 12min: 24, 15min: 22, 18min: 22); (LPA_5_) N = 1, n = 10 (for each time point); (Bb) N = 1, n = (3 for each time point); (LPA_50_) N = 7, n = (1min: 32, 2min: 32, 3min: 33, 6min: 21, 9min: 21, 12min: 21, 15min: 21, 18min: 20) **(d)** Cell-cell contact size (A_c_) at 12 min contact time for control doublets and doublets exposed to Bb (10 μM) or different amounts of LPA (5 - 50 nM) plotted against cortical tension (T_c_) values measured by AFM. Error bars denote standard deviation. For T_c_ measurements: N = 3 and (single cell Ctrl) n = 287; (single cell Bb) n = 88; (single cell LPA_5_) n =142; (single cell LPA_50_) n = 294. For A_c_ measurements: N and n as in Fig. 1c (time point 12min). **(e)** A_c_ in doublets exposed to 50 nM LPA as a function of time in culture before and after adding 10 μM Bb to the culture medium. Grey area denotes standard deviation; N = 3, n = 16. If not stated otherwise, N corresponds to number of experiments and n to number of cell doublets.

To systematically investigate how changes in cortical tension affect contact expansion in doublets, we determined how global cortical tension is altered in progenitor cells upon exposure to Bb or different concentrations of LPA by employing Atomic Force Microscopy (AFM)^29,30^, and how those changes relate to contact expansion. We found that A_c_ was maximized when cortical tension was left unaltered, while it dropped when cortical tension was either elevated in the presence of LPA (5-50 nM) or diminished upon exposure to Bb (10 μM; Fig. 1d). This suggests that the threshold level or cortical tension delineating the transition point from where on cortical tension is not promoting but inhibiting contact expansion is close to the cortical tension level of untreated progenitor cells. To exclude the possibility that the effect of LPA on cell-cell contact expansion is not due to its activity in elevating cortical contractility, but rather by modulating other potential effector processes, such as actin polymerization^31^, we reduced Myosin II activity in LPA-treated progenitor cell doublets and determined how this affects the ability of LPA in restricting contact expansion. While cell-cell contact expansion was strongly restricted in cell doublets exposed to 50 nM LPA, this restriction was abrogated when 10 μM Bb was also added to the culture medium (Fig. 1e). This confirms that LPA restricts cell-cell contact expansion by elevating cortical actomyosin contractility. Collectively, our findings so far suggest that cortical tension has a dual function in controlling contact expansion with low to normal levels of cortical tension promoting and high levels of cortical tension inhibiting contact expansion.

To explore the mechanisms by which high cortical tension limits contact expansion, we turned to previous observations that cortical tension might increase Cadherin adhesion complex clustering and stability at cell-cell contact sites^16^. To visualize Cadherin complex dynamics in progenitor cell doublets, we took advantage of a transgenic line, which expresses a Citrine-tagged version of α-ECatenin (Ctnna1) under its endogenous promoter (Tg*(ctnna-citrine)^ct3a^)* colocalizing with E-cadherin/Cdh1 (Fig. S2), the prime classical cadherin mediating zebrafish germ layer progenitor cell-cell contact formation^29^. Analyzing the subcellular distribution of Ctnna1 at the cell-cell contact in control and LPA-treated doublets (50 nM) revealed a stronger accumulation of Ctnna1 at the contact edge in treated compared to control doublets (Fig. 2a). This enhanced accumulation of Ctnna1 at the contact edge was accompanied by similar changes in cortical Actin localization (Fig. 2a). Myosin II, in contrast, did not show any detectable accumulation at the contact edge in both treated and control doublets (Fig. 2a). High-resolution analysis of Ctnna1 distribution at the contact further showed both brighter and larger Ctnna1 clusters in LPA-treated versus control doublets (Fig. 2b), which colocalized with E-cadherin/Cdh1 (Fig. S2). Together, this suggests that elevating cortical tension promotes E-cadherin/Ctnna1 adhesion complex accumulation and clustering at the contact edge.

**Figure 2.**
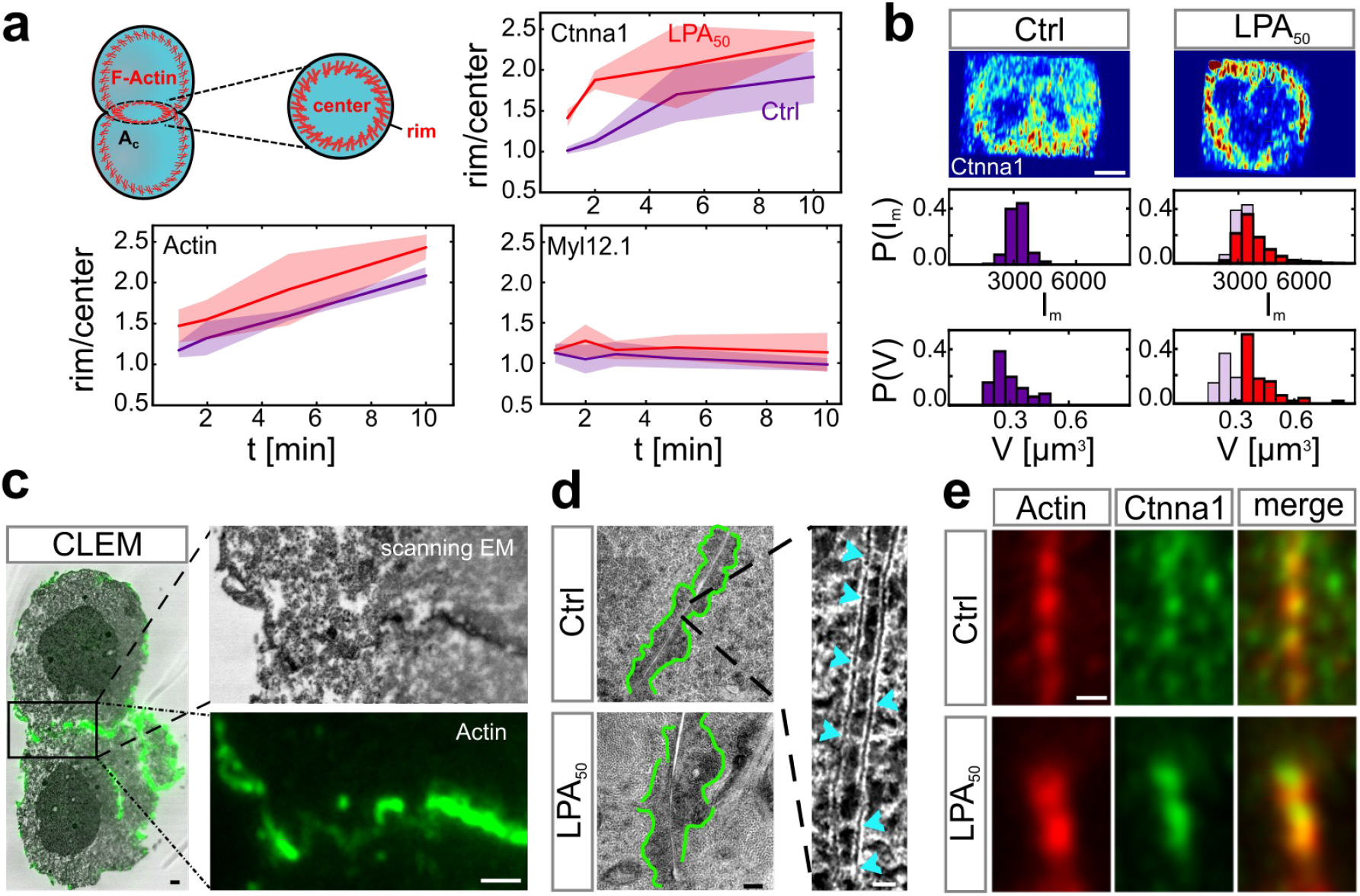
Cortical tension triggers Ctnna1/Actin clustering at the contact of cell doublets. **(a)** Rim to centre intensity ratios of core components of the Cadherin-adhesion complex in doublets. Schematic on the left shows the rim and centre regions of the cell-cell contact where the fluorescence mean intensities were measured in control doublets (red line) and doublets exposed to LPA 50 nM (blue line). Doublets were fixed and analysed for each time point separately (1, 2, 5, 10 min contact time). F-actin was visualized by Phalloidin with N = 3 and (Ctrl) n = 5, 3, 3, 3 (corresponding to the different contact times mentioned above); (LPA) n = 6, 5, 4, 5. Ctnna1 was visualized by immunohistochemistry with N = 3 and (Ctrl) n = 5, 3, 5, 5; (LPA) n = 4, 3, 4, 3. Myosin II was visualized by Myl12.1-eGFP expression with N = 1 and (Ctrl) n = 9, 8, 7, 8; (LPA) n = 5, 5, 4, 2. Shadowed area denotes standard deviation. **(b)** Exemplary sub-diffraction limited confocal images of Ctnna1 subcellular distribution at the cell-cell contact of control doublets (left) and doublets exposed to 50 nM LPA (right). Quantifications below show cluster mean intensity (I_m_) and volumes (V) of the 50 largest clusters of each cell-cell contact. Blue shadows in the right panel denote control conditions. (Ctrl) N = 1, n = 3; (LPA) N=1, n = 5. **(c)** Correlative light and electron microscopy (CLEM) images with F-actin visualized by Phalloidin-Alexa 488 (green). Right panels are zoomed-in images of the boxed region in the left panel. Scale bars, 1 μm. **(d)** Electron microscopy (EM) images of electron-dense clusters (outlined with green) at cell-cell contacts in control doublets (upper panel) and doublets exposed to 50 nM LPA (lower panel). Scale bar, 200 nm. Right image is a zoomed-in image of Cadherin-like clusters with individual clusters depicted by light blue arrowheads. Scale bar, 20 nm. **(e)** Representative ‘Airy Scan’ images of F-actin (red) and Ctnna1 (green) co-localizing in clusters at the cell-cell contact of control doublets (upper panel) and doublets exposed to 50 nM LPA (lower panel). Cell doublets were fixed after 30 min contact time and visualized by Phalloidin (F-actin) and immunohistochemistry (Ctnna1). Scale bar, 10 μm. If not stated otherwise, N corresponds to number of experiments and n to number of cell doublets.

To analyze how cortical tension affects Ctnna1 adhesion clusters and their association to the Actin cytoskeleton at high resolution, we performed correlative light and electron microscopy (CLEM) on E-cadherin clusters in control and LPA-treated doublets (Fig. 2c). By labelling F-actin with Phalloidin, we were able to determine how putative E-cadherin adhesion complexes were associated with the Actin cytoskeleton. Consistent with our high-resolution light microscopy analysis (Fig. 2b), we found junctional structures closely resembling adherens junctions at the contact of both control and LPA-treated doublets (Fig. 2c,d). Moreover, we observed Actin accumulations adjacent to those structures, which appeared enlarged in LPA-treated doublets (Fig. 2d), suggesting that cortical tension might not only increase the size of E-cadherin clusters but also of the adjacent Actin accumulations. To further test this notion, we performed high-resolution fluorescence microscopy of contacts stained for both Ctnna1 and F-actin in control and LPA-treated doublets, revealing larger Actin accumulations adjacent to larger Ctnna1 clusters in LPA-treated compared to control doublets (Fig. 2e).

Tension-induced Cadherin cluster formation at cell-cell contacts has previously been associated with reduced adhesion molecule turnover^16^. To test whether adhesion complex turnover at germ layer progenitor cell-cell contacts is affected by changes in cortical tension, we performed Fluorescence Recovery After Photobleaching (FRAP) experiments of Ctnna1 in control and LPA-treated doublets (Fig. 3a). Interestingly, by using kymographs to outline spatiotemporal changes in Ctnna1 intensity after photobleaching (Fig. 3b) and quantifying those changes as a function of time (Fig. 3c), we found that within seconds - the timescale relevant for LPA affecting contact expansion in progenitor cell doublets (Fig. 1c) - recovery of Ctnna1 from adjacent non-bleached areas of the contact edge was strongly slowed down in progenitor cell doublets exposed to 50 nM LPA (Fig. 3c). This suggests that increasing cortical tension not only increases Cadherin clustering, but also reduces Cadherin adhesion complex turnover at the contact site.

**Figure 3.**
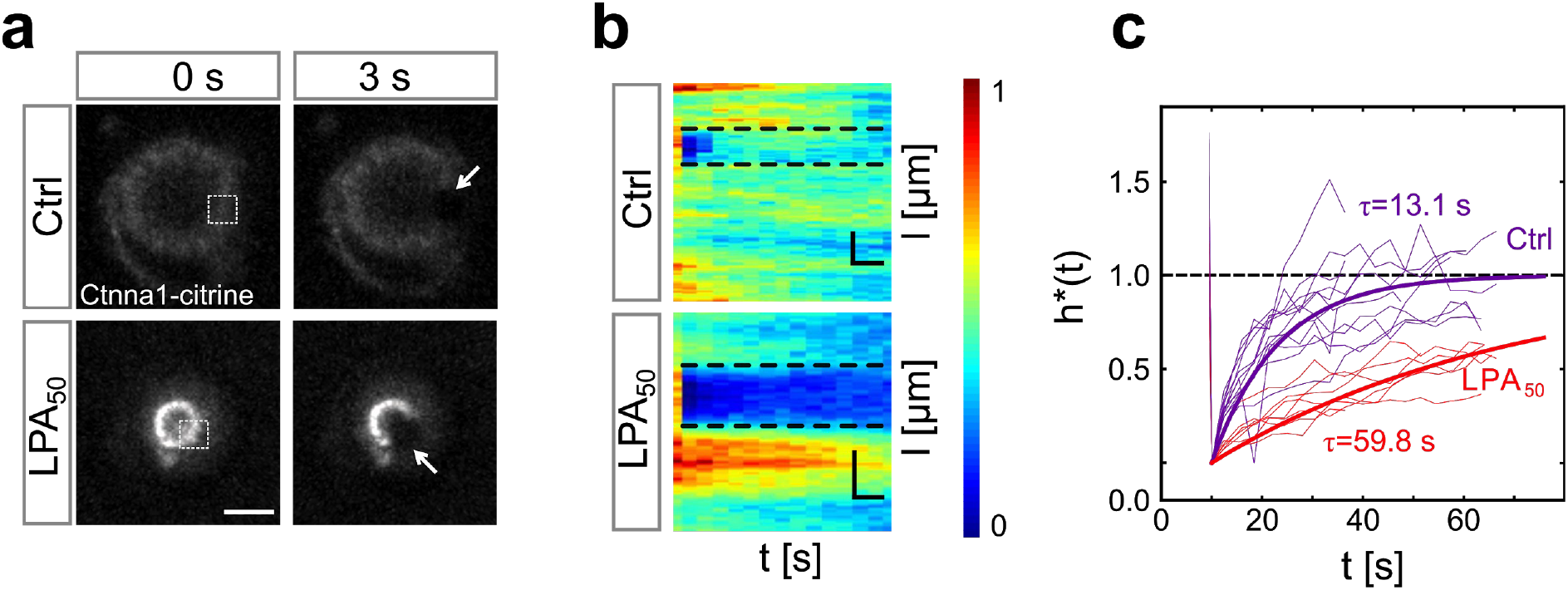
Cortical tension reduces Ctnna1 turnover at the contact of cell doublets. (a) FRAP analysis of Ctnna1 turnover at the contact of progenitor cell doublets. Fluorescence images of Ctnna1 localization within the contact plane of control doublets (upper row) and doublets exposed to 50 nM LPA (lower row) in the last pre-bleach (left panels) and first post-bleach frames (right panels). Boxed regions (left column) and arrows (right column) outline bleached regions. Scale bar, 10 μm. **(b)** Normalized intensity kymographs of Ctnna1 recovery after photobleaching at the cell-cell contact edge of control doublets and doublets exposed to 50 nM LPA. Scale bars, 6s (horizontal) and 10 μm (vertical). **(c)** Quantification of Ctnna1 fluorescence intensity within the bleached regions at the contact edge of control doublets (purple) and doublets exposed to 50 nM LPA (red) as a function of time after photobleaching. τ denotes recovery characteristic time scale. Thin lines denote individual cases and thick lines averages. (Ctrl) N = 3, n = 12; (LPA_50_) N = 2, n = 7. See Methods section for details. If not stated otherwise, N corresponds to number of experiments and n to number of cell doublets.

Next we asked how cortical tension promotes E-cadherin clustering and stability. Previous studies have suggested that cortical actomyosin tension transduced to the Cadherin adhesion complex promotes anchoring of the complex to the cortex, and that increased cortical anchoring enhances Cadherin stability and clustering^32,33,16^. For testing whether such mechanism can also explain the observed effect of cortical tension on E-cadherin clustering and turnover in germ layer progenitor cell doublets, we first determined how tension within the E-cadherin adhesion complex is changed upon alterations in cortical actomyosin tension. For visualizing tension within the E-cadherin adhesion complex, we took advantage of previous observations that tension at Cadherin adhesion sites leads to unfolding of α-Catenin and, as a result of this, enhanced recruitment of Vinculin to those sites^34,7,35,20^. Determining the amount of Vinculin at the contact edge in control versus LPA-treated progenitor cell doublets revealed strongly increased Vinculin accumulation upon exposure to LPA (50 nM) already 1 min after contact initiation (Fig. S3a,b), suggesting that LPA increases tension within the cadherin adhesion complex.

Next we asked how increased cortical tension and thus tension within the Cadherin adhesion complex affects anchoring of the adhesion complex to the actomyosin cortex. To this end, we used a dual micropipette aspiration assay (DPA) to determine the de-adhesion strength of control and LPA-treated progenitor cell doublets^1^. We have previously shown that when separating progenitor cell doublets using DPA, the E-cadherin adhesion complex at the cell-cell contact ruptures first at its linkage to the actomyosin cortex, suggesting that the de-adhesion force is limited by the cortical anchoring strength of the E-cadherin adhesion complex and thus can be used to determine cytoskeletal anchoring of E-cadherin in those cells^1^. Comparing the de-adhesion force of progenitor cell doublets in the presence or absence of LPA revealed higher de-adhesion forces when cell cortex tension was elevated in cell doublets by exposing them to 50 nM LPA (Fig. 4a). Given that the total amount of clustered and unclustered E-cadherin at the contact was not increased upon LPA treatment (as measured by determining the integrated intensity of Ctnna-citrine at the cell-cell contact with Ctrl = 7.5 +/−2.8 a.u. [N=3, n=2, 6, 6] and LPA = 4.8 +/−1.6 a.u. [N=3, n=1, 6, 2]; see also Fig. 2a,b), this suggests that cortical tension promotes anchoring of the E-cadherin to the actomyosin cortex in progenitor cell doublets.

**Figure 4.**
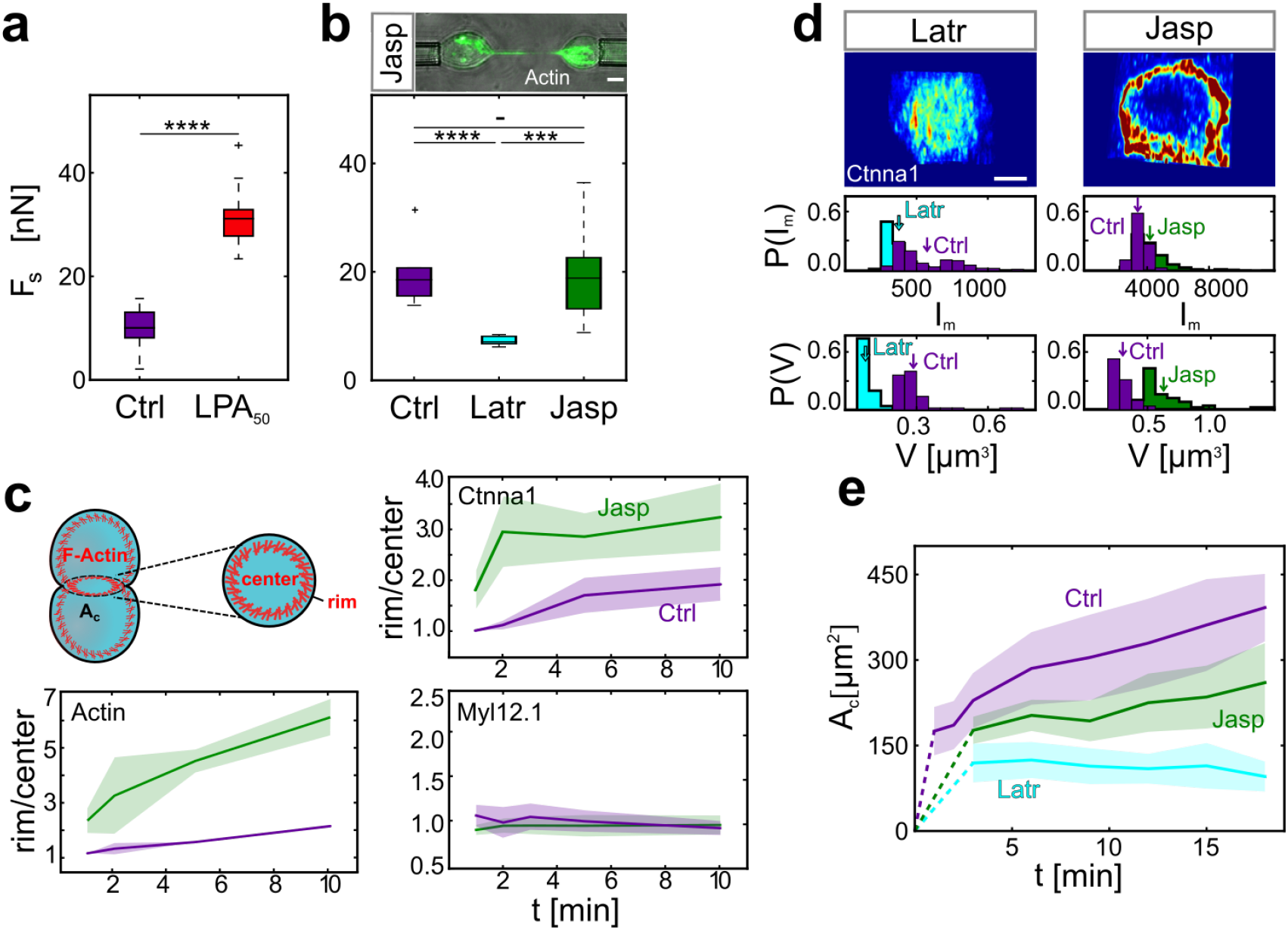
Enhanced cytoskeletal anchoring of the Cadherin adhesion complex by cortical tension limits contact expansion in doublets. **(a)** De-adhesion forces (F_s_) for control progenitor cell doublets and doublets exposed to 50 nM LPA after 10 min contact time. (Ctrl) N = 13, total n = 17; (LPA_50_) N = 7, total n = 12. **** *P*-value = 7.2e-12, Student *t*-test. **(b)** F_s_ for control doublets and doublets exposed to 300 nM Latr or 100 nM Jasp. Upper panel shows Actin-rich tethers formed between the cells during separation in the presence of Jasp. (Ctrl) N = 3, n = 10; (Latr) N = 1, n = 8; (Jasp) N = 3, n = 13. *P*-value **** (Ctrl-Latr) 1.92e-5, *** (Latr-Jasp) 6.96e-4, ns (Ctrl-Jasp) not significant; *t*-test with Bonferroni correction for multiple comparisons. **(c)** Rim to centre mean intensity ratios for F-actin, Myosin II, and Ctnna1 as a function of contact time (1, 2, 5, 10 min) in the presence or absence of Jasp. F-actin was visualized by Phalloidin with N = 3 and (Ctrl) n = 5, 3, 3, 3 (corresponding to the different contact times mentioned above); (Jasp) n = 3, 4, 3, 4. Ctnna1 was visulaized by immunohistochemistry with N = 1 and (Ctrl) n = 5, 3, 5, 5; (Jasp) n = 3, 3, 3, 4. Myosin II was visualized by Myl12.1-eGFP expression with N = 1 and (Ctrl) n = 9, 8, 7, 8; (Jasp) n = 12, 9, 11, 8. Shadowed area denotes standard deviation. **(d)** Exemplary **‘** Airy Scan’ images of Ctnna1 subcellular localization at the contact edge of doublets exposed to Latr or Jasp. Quantifications below show cluster mean intensity (I_m_) and volumes (V) of the 50 largest clusters of each cell-cell contact. Arrows indicate distribution means. N =1 and (Latr) n = 7; (Jasp) n = 5. **(e)** Cell-cell contact size (A_c_) of control doublets and doublets exposed to Jasp or Latr as a function of contact time. Dotted lines connect contact formation (0 min) with the first time point when data were collected. Shadowed area denotes standard deviation with (Ctrl) N and n as in Fig. 1c; (Jasp) N = 4, n = (3min: 6, 6min: 6, 9min: 16, 12min: 16, 15min: 16, 18min: 16); (Latr) N = 4, n = (3min: 21, 6min: 21, 9min: 21, 12min: 21, 15min: 21, 18min: 9). If not stated otherwise, N corresponds to number of experiments and n to number of cell doublets.

To determine whether increased cortical anchoring of the Cadherin adhesion complex by tension enhances Cadherin clustering at cell-cell contacts, we analyzed how changes in cortical anchoring affect E-cadherin clustering in progenitor cell doublets. For modulating the anchoring strength of the E-cadherin to the actomyosin cortex in these cells, we sought to interfere with F-actin network stability rather than changing specific cadherin adhesion complex components, whose anchoring function is not yet fully understood^36^. To modulate cortical Actin network stability, we exposed doublets to Latrunculin (Latr), blocking Actin polymerization and thus destabilizing the cortical Actin network, or Jasplakinolide (Jasp), promoting actin polymerization and network stability^33,16^. First, we analyzed how exposure to Latr and Jasp affects cortical anchoring of the E-cadherin complex by determining the de-adhesion force of progenitor cell doublets as a readout of the cortical anchoring strength in the presence of Latr or Jasp. The de-adhesion force of cell doublets was decreased when F-actin was destabilized with Latr (Fig. 4b), suggesting that exposure to Latr reduces the cortical anchoring strength of the E-cadherin adhesion complex. Conversely, when progenitor cell doublets were exposed to 100 nM Jasp to stabilize the F-actin network, the apparent de-adhesion force remained largely unchanged (Fig. 4b). However, multiple Actin-containing tethers were typically observed between the separating cells (Fig. 4b), suggesting that contacts were not fully separated. While these measurements did not reveal the complete separation force of Jasp-treated doublets, they show that the cortical anchoring strength of E-cadherin in the presence of Jasp exceeds the resistance of the Actin cytoskeleton to considerable deformation when pulled into tethers. Importantly, the formation of Actin-filled tethers upon separation was not observed when increasing cytoskeletal anchoring of the E-cadherin adhesion complex by LPA^1^, presumably as a result of LPA, but not Jasp, also promoting cortical actomyosin tension resisting actin network deformation.

We then asked how increasing the cortical anchoring strength of the E-cadherin adhesion complex would affect E-cadherin clustering in cell doublets. Analyzing the distribution of Ctnna1 at the contact in Jasp-treated doublets revealed enhanced accumulation at the contact edge (Fig. 4c), which was accompanied by similar changes in Actin localization (Fig. 4c). Myosin II distribution, in contrast, did not show such changes in response to Jasp treatment (Fig. 4c). Furthermore, high-resolution analysis of Ctnna1 clustering at the contact edge in Jasp-treated doublets revealed brighter and larger Ctnna1 clusters when the anchoring strength was elevated upon exposure to Jasp (Fig. 4d).

Finally, we asked how modulating E-cadherin complex anchoring to the cortex and, consequently, E-cadherin clustering at cell-cell contacts would affect cell-cell contact expansion in doublets. To this end, we analyzed contact expansion in doublets exposed to Latr or Jasp, decreasing and increasing anchoring strength, respectively. Strikingly, we found that contact expansion in doublets was strongly reduced not only upon Actin destabilization via Latr but also by stabilizing it in the presence of Jasp (Fig. 4e). Together, these findings suggest that cortical tension limits contact expansion by enhancing the anchoring of the Cadherin adhesion complex to the actomyosin cortex and, as a result of this, Cadherin clustering and stability at cell-cell contacts.

To directly challenge this conclusion, we sought to test how disrupting cortical anchoring of the Cadherin adhesion complex interferes with the ability of high cortical tension in restricting contact expansion in doublets. To this end, we substituted the endogenous E-cadherin in progenitors forming doublets with a truncated version of N-cadherin lacking its cytoplasmic tail (Cdh2Δcyto) and thus its ability to anchor to the cortical actomyosin network^1^. We used N-cadherin (Cdh2) instead of E-cadherin as expressing sufficient amounts of exogenous Chd2 to substitute for the function of endogenous E-cadherin in germ layer progenitors turned out to be easier than expressing exogenous E-cadherin^1^. We first tested whether the cytoplasmic tail of Cdh2 is required for LPA to promote Cadherin-anchoring to the Actin cortex. When comparing the de-adhesion forces of doublets in the presence versus absence of LPA expressing either full-length Cdh2 (Cdh2FL) or its truncated version (Cdh2Δcyto), we found that LPA increases the de-adhesion force only in doublets expressing Cdh2FL but not Cdh2Δcyto (Fig. 5a). This suggests that the cytoplasmic tail of Cdh2 is required for LPA enhancing Cdh2 cytoskeletal anchoring. To further test whether the cytoskeletal anchoring of Cdh2 is needed for LPA restricting contact expansion in doublets, we compared contact expansion in cell doublets expressing either Cdh2FL or Cdh2Δcyto. Strikingly, doublets expressing Cdh2Δcyto failed to show any recognizable changes in contact expansion when exposed to LPA (Fig. 5b,c), while doublets expressing Cdh2FL displayed similar changes in contact expansion and size upon LPA treatment (50 nM) than found in control cell doublets (Fig. 5b,c). Collectively, these findings support the notion that cortical tension restricts contact expansion by promoting the cytoskeletal anchoring of the Cadherin adhesion complex.

**Figure 5.**
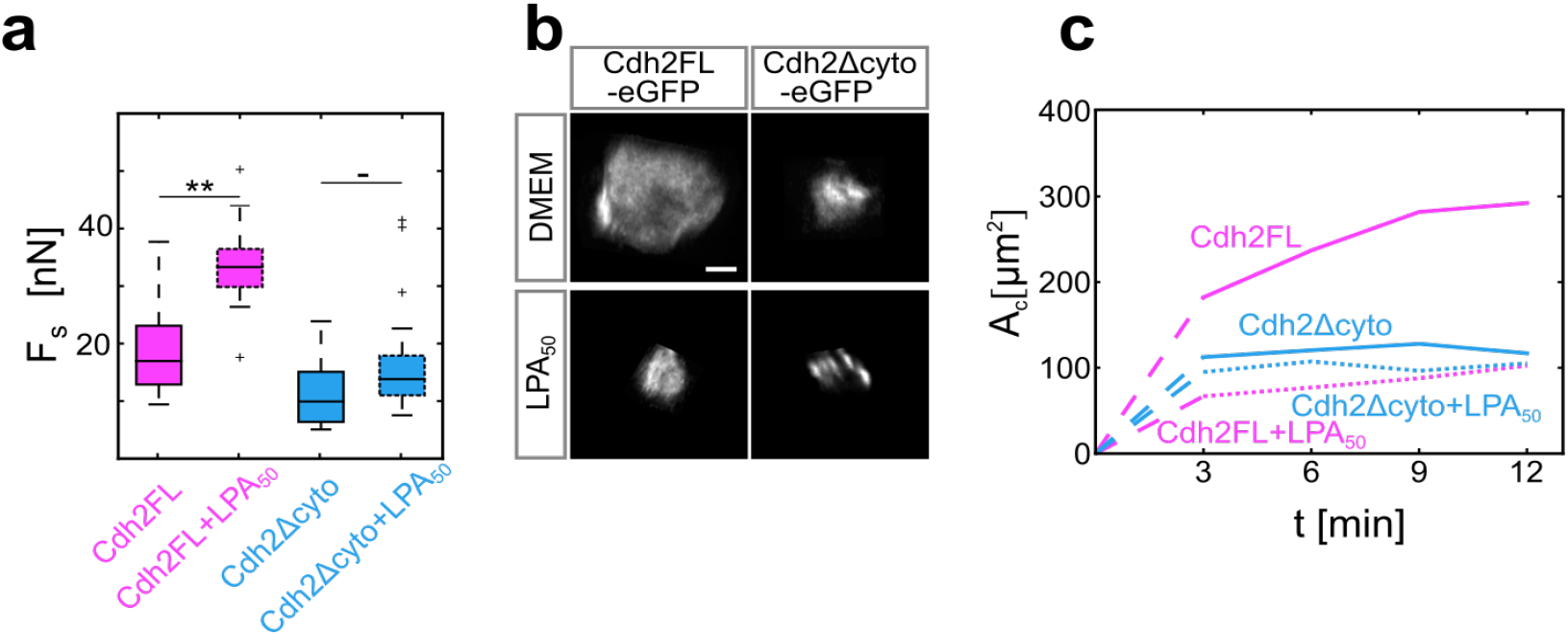
Defective cytoskeletal anchoring of the Cadherin adhesion complex suppresses the effect of cortical tension on contact expansion. **(a)** De-adhesion forces (F_s_) in doublets expressing either full-length Cdh2 (Cdh2FL, pink) or Cdh2 lacking its cytoplasmic C-terminus (Cdh2Δcyto, blue) in the presence (dashed boxes) or absence (solid boxes) of 50 nM LPA. (Cdh2FL) N = 3,n=10; (Cdh2FL+LPA_50_) N = 3, n = 12; (Cdh2DeltaCyto) N = 9, n = 30; (Cdh2DeltaCyto+LPA_50_) N = 7, n = 20. *P*-values: ** (Cdh2FL-Cdh2FL+LPA_50_) 9.9e-3, ns (Cdh2DeltaCyto-Cdh2DeltaCyto+LPA_50_) not significant; *t*-test with Bonferroni correction for multiple comparisons **(b)** Subcellular localisation of Cdh2FL-eGFP and Cdh2Δcyto-eGFP at the contact of doublets in the presence or absence of 50 nM LPA. Scale bar, 5 μm. **(c)** Cell-cell contact size (A_c_) in doublets expressing either Cdh2FL (pink) or Cdh2Δcyto (blue) in the presence (dashed line) or absence (solid line) of 50 nM LPA. Dotted lines connect contact formation (0 min) with the first time point when data were collected. Standard deviations are shown in Fig S4. (Cdh2FL) N = 2, n = (3min: 8, 6min: 8, 9min: 8, 12min: 8); (Cdh2Δcyto) N = 2, n = (3min: 14, 6min: 14, 9min: 14, 12min: 14); (Cdh2Δcyto+LPA_50_) N = 2, n = (3min: 9, 6min: 9, 9min: 9, 12min: 9); (Cdh2FL+LPA_50_) N = 2, n = (3min: 10, 6min: 10, 9min: 10, 12min: 10). If not stated otherwise, N corresponds to number of experiments and n to number of cell doublets.

## Discussion

Our findings suggest a non-monotonic relationship between cell cortex tension and cell-cell contact size. At low and moderate levels of cortical tension, tension positively scales with contact size, consistent with a simple model where the level of cortical tension pulling on the contact edge determines the size of the contact once force equilibrium between the contacting cells is reached. At high levels of cortical tension, however, the contact size inversely scales with the level of tension. Importantly, this does not argue against the general concept of force equilibrium between the contacting cells determining contact size, but rather suggests that the ability of the contact to expand might be force-sensitive, permitting fast contact expansion at low to moderate levels of cortical tension, while considerably slowing it down at high tension levels.

We also show that cortical tension diminishes the ability of the contact to expand by increasing E-cadherin anchoring to the cortical actomyosin cytoskeleton, and that this enhanced anchoring leads to E-cadherin clustering and reduced turnover of the E-cadherin adhesion complex at the contact. Previous studies have provided evidence that unfolding of α-catenin and stabilization of the Actin network are involved in mediating the effect of tension on E-cadherin clustering by promoting cytoskeletal anchoring of the cadherin adhesion complex^21,38,35,20,39,16^. Our observations suggest that cortical tension not only triggers those mechanosensitive effector processes, but that activation of those processes has severe morphogenetic consequences by restricting contact expansion and enhancing contact strength.

Commonly, contact size is assumed to scale with contact strength, and there are multiple cases where such a relationship has been documented^1,40,12^. Our findings of an inverse relationship between contact size and strength point at the possibility that some processes might benefit from cell-cell contacts being simultaneously small and strong. For instance, during collective or chain cell migration, cells need to establish stable contacts with their neighbors and, at the same time, retain contact-free interfaces that allow them to form protrusions required for cell migration^41,42^. This points at the possibility that the non-monotonic co-regulation of contact size and strength as a function of cortical tension might reflect specific features of these contacts: at low to moderate levels of cortical tension, contact size might be less important as contacts are likely to be more transient and flexible. At high cortical tension, in contrast, contacts are expected to be long-lived and stable, and thus contact size will more permanently affect other processes requiring cell-cell contact free interfaces, such as cell protrusion formation and cell-matrix adhesion. Whether and how the combined effect of contact size and strength affects specific biological processes remains to be investigated.

There is manifold evidence for cadherin-mediated cell-cell contacts being mechanosensitive^43,44^. While the molecular and cellular mechanisms underlying this mechanosensitivity have been studied in detail^1,45,46^, comparably little is yet known about its function in contact formation and maintenance. Our findings of an important function for mechanosensation at cadherin-mediated cell-cell contacts in controlling contact expansion identify a yet unknown role of mechanosensation in determining contact dynamics and strength.

## Materials and Methods

### Fish lines and husbandry

Zebrafish maintenance was carried out as described^47^. Embryos were grown at 28-31°C in E3 embryo medium (5 mM NaCl, 0.17 mM KCl, 0.33 mM CaCl, 20.33 mM MgSO_4_) and prior to experiments dechorionated in Danieau’s embryo medium (58 mM NaCl, 700 μM KCl, 400 μM MgSO_4_, 600 μM Ca(NO_3_)_2_ and 5 mM Hepes at pH 7.2). Embryos were staged based on morphological criteria^48^. The following fish-lines were used: wild-type embryos were obtained from the Tup-Longfin (TL) and AB background; transgenic fish-lines used were Tg(*ctnna-citrine*)^*ct3a*^ ^49^, Tg(*bAct:myl12.1-eGFP*)^1^, Tg(*bAct:myl12.1-mCherry*)^1^, Tg(*bAct:LifeAct-eGFP*)^50^, and Tg(*actb1:mCherry–utrCH*)^50^.

### Cell culture

To prepare primary cultures of zebrafish progenitor cells for live imaging, embryos consisting of ectoderm cells only (see also ‘mRNA and *morpholino* injections’ below) were raised as described above. After embryo dechorionation in Danieu’s medium at sphere stage (4hpf) the blastoderm cap was dissected from the yolk cell using forceps and transferred first to CO_2_ - independent DMEM/F12 culture medium (DMEM/F12, Invitrogen, complemented with L--Glut, 15 mM Hepes and 100 U/mL penicillin plus streptomycin, adjusted at pH 7.5, sterilized using 0.45 μm pore filters, and preheated to 28° C) and then to an Eppendorf tube containing 200 μl culture medium using a glass pipette. Blastoderm caps were afterwards mechanically dissociated into single cells by gentle tapping on the tube, and the cells were then transferred into a fetal bovine serum (FBS, Gibco) coated glass bottom dish (MatTek glass bottom dish, 35 mm, MatTek Corporation). Cell-cell contact formation was initiated by gently bringing two cells together using micropipettes^1,2^, and the newly formed cell doublet was imaged up to 20 min.

### mRNA and *morpholino* injections

Zebrafish embryos were induced to consist of ectoderm progenitors only by micro-injection of one-cell stage embryos with 100 pg *lefty1* mRNA. To substitute endogenous E-cadherin with controlled amounts of full-length Cdh2FL or C-terminal truncated Cdh2Δcyto, 8 ng E-cadherin/*cdh1 morpholino* (MO) (5′- TAAATCGCAGCTCTTCCTTCCAACG −3′, GeneTools) together with 100 pg of *cdh2FL-eGFP* or 100 pg *cdh2Δcyto-eGFP* mRNA^51,1^ were injected into one-cell stage embryos. To increase cortical tension, 5 pg of *ca RhoA^28^*, and to visualize subcellular Vinculin distribution, 150 pg of *vinculinB-eGFP* were injected into one-cell stage embryos. Synthetic mRNA was produced by using the SP6 mMesage mMachine kit (Ambion).

### Immunostaining

Single progenitor cells were obtained as described in ‘Cell culture’ above and allowed to seed on MatTek dishes for 30 min. Cells were then fixed with 4% paraformaldehyde (PFA, Sigma-Aldrich) in DMEM/F12 for 10 min at room temperature (RT), washed 3x with phosphate buffered saline (PBS, Sigma-Aldrich) to remove the PFA, and incubated in PBS with 0.3% Triton X100 (Merck) (PBT) for 30 min at RT to permeabilize the plasma membrane. PBT was subsequently replaced with blocking solution consisting of PBT with 1% DMSO (Sigma-Aldrich) and 10% goat serum (GS, Gibco) for 1 h at RT before primary antibodies diluted in blocking solution were added overnight at 4°C. Cells were washed 3x with PBS at RT, and secondary antibodies diluted in blocking solution were added for 2 h at RT, followed by three washes with PBS at RT to remove the antibodies. The following primary antibodies were used: αE-catenin (1:1000, Sigma-Aldrich C2081), Vinculin 1:100 (1:100, Sigma-Aldrich V4505) and E-cadherin (1:250, MPI-CBG^52^). As secondary antibodies, fluorescently Alexa-488, Alexa-647 or Alexa-568 coupled secondary antibodies (1:250, Molecular Probes) were used. For labelling F-actin, Phalloidin (1:250, Invitrogen) was used. Immuno-labelled cells were imaged on a Zeiss Observer inverted microscope equipped with a Spinning Disc System (see also ‘Imaging’ below).

### Imaging acquisition

Fluorescence imaging of cells was performed on the Spinning Disc System (Andor Revolution Imaging System; Yokogawa CSU-X1) placed on an inverted microscope (Axio Observer Z1 Zeiss) using a 40x/1.2 NA water immersion lens (Zeiss) for time-lapse imaging and a 100x/1.4 NA Oil (Zeiss) for still images. The setup was equipped with a motorized piezo stage, stage heating and objective heater units. Single and dual color fluorescence images were acquired using 488 nm and 561 nm laser lines with an optical slicing of 0.5 μm. 30 mW maximum laser output power was used, and images were acquired using an iXon DU-897-BV EMCCD camera (Andor Technology) with exposure times set to 100-300 ms and frame rates between 1-2 s. Resulting image z-stacks were rotated using Imaris 9.1.2 (BitplaneTM) to obtain cross sections of cell-cell contacts between cell doublets. High-resolution images of endogenous E-cadherin/Cdh1, Ctnna1 and Actin clusters were obtained using a Inverted Zeiss LSM 880 confocal / ‘Airy Scan’ using a 63x/1.4 NA Oil (Zeiss), and image analysis was performed using ImageJ software^53^.

### Image analysis

Visualisation of Ctnna1 clusters was performed using deconvolved (Zeiss ZEN 2.3) z-stacks of ‘Airy Scan’ confocal images of cell doublets, with 50–100 images per stack and 0.19 μm z-increment. Protein cluster volume and fluorescence intensity were detected and quantified using Imaris 9.1.2 (BitplaneTM)^54^.

VinculinB-GFP fluorescence intensity was quantified from z-stack images of cell-cell contacts by first selecting a plane in the stack, where the cell-cell contact appeared the largest (corresponding to the middle of the contact), and then measuring average intensity in two 3×3 pixel regions located at the edges of the cell-cell contact (rim intensity) and two in the middle of the contact line (centre intensity). From that, rim to centre ratio was calculated.

### Fluorescence Recovery After Photobleaching (FRAP) analysis

For each experiment, two progenitor cells from Tg(*ctnna-citrine*)^*ct3a*^ line were placed in a polymer-well mounted on a MatTek dish using pipettes and allowed to form a contact. Subsequently, the cell-cell contact plane was aligned with the focal plane. After cells had been in contact for 10+/−2 min, at least 10 frames of pre-bleach fluorescence intensity were recorded and then a 488 laser (300 pulses, 300 ms, dwell time 30 μs) was used to bleach a small rectangular area on the contact edge (region of interest, ROI) with the size of 8×8 pixels. Imaging of the bleached area was performed at 3.3 frames/s with an image size of 512×512 pixels for at least 60 s before the contact plane would typically drift out of the focal plane.

For each bleached contact a polar transformation was performed around the centre of the contact for a pre-bleach image followed by a post-bleach image series (300 ms from the recorded time series) using ImageJ Polar Transformer plugin (https://imagej.nih.gov/ij/plugins/polar-transformer.html). In the transformed images a line profile was taken along the contact edge (line thickness 9 pixels), and the radial span of the bleached region was recorded. Subsequent stacks were aligned into kymographs using a cross-correlation method to correct for cell doublet rotation. The intensity in the bleached region *I_b_* (*t*) and outside of it *I_u_* (*t*) was then recorded over time. To correct for acquisition photobleaching, the following transformations were applied to the data:

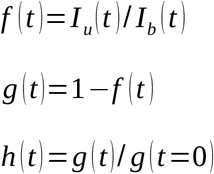

where t=0 corresponds to the first time point after bleaching. In this way *h* (*t*) *→* 0 corresponds to full recovery of the signal. Single exponential recovery equations were then fitted using nonlinear least squares to *h* (*t*)=*e*^−*τ*∗*t*^, where *τ* is the characteristic recovery time. Fitting errors were calculated as squares of diagonal elements of the covariance matrix. In Fig. 2c *h*^❑^(*t*)=1−1/ *h* (*t*) is shown to conform to typical representation of the FRAP data.

### Rim to centre ratio and contact size analysis

A number of freshly formed cell doublets (typically 1-4) expressing Citrine-tagged Ctnna1 were followed for up to 20 min (at which point the cells would typically drift from the field of view or divide) and imaged every 3 min on a spinning-disk microscope using z-stacks. From each doublet 3 sub-stacks were cut out for every time point: one containing the cell-cell contact with thickness *d* and two stacks containing small cytosol volumes from each cell, from which the average cytosolic signal intensity (*I_c_*) was calculated. Subsequently, the cell-cell contact was projected on a plane (sum of the signal) and background signal *I_c_*∗*d* was subtracted from it. The contour of the contact was detected by applying a combination of thresholding (typically at the 1.5-2.5 times background intensity, visually assessed to contain whole cell-cell contacts) and subsequent dilation of the binarized image to account for uneven distribution of adhesion molecules at the cell-cell contact rim. An ellipse was then fitted to the contour using a set of custom-made scripts in order to calculate the surface area *C_c_*=*π*∗*a*∗*b,* where *a* and *b* denote ellipse semi-axes. Due to the dotted and discontinuous nature of GFP signal in cells expressing Cdh2FL and Cdh2DeltaCyto the automatic contact size measurement was not possible. In these cases a diameter of each cell-cell contact was manually measured using ImageJ.

### Integrated intensity analysis

Cell doublets were placed in polymer wells as described above, with cell-cell contacts selected that had the contact plane well aligned with the imaging plane. Background values were taken as average fluorescence intensity in a neighbouring empty polymer well and subtracted from each image. Cell-cell contacts were outlined and the sum of the raw fluorescence intensity was calculated.

### Dual Pipette Assay (DPA)

Single progenitor cells were prepared as described in ‘Cell culture’ section above. MatTek glass-bottom dishes were passivated by incubation in heat-passivated FBS (Invitrogen) for at least 20 min at RT. Glass pipettes with a diameter of 8 μm (Biomedical Instruments) were passivated in the same way for 7 min, washed with PBS and connected to a Microfluidic Flow Control System (Fluigent, Fluiwell) with negative pressure range of 7 - 750 Pa, accuracy of 7 Pa and change rate of 200 Pa/s on two independent channels. Micropipette movement was performed by micromanipulators (Eppendorf, Transferman Nk2), which together with the pressure were controlled via a custom-programmed Labview (National Instruments) interface. Dissociated cells from 1-2 embryos were transferred to the MatTek dishes in 4 ml DMEM/F12 and allowed to seed for at least 10 min. To manipulate single cells, ~ 20 Pa negative pressure in the pipettes was used.

For each measurement, two healthy looking cells were selected, put in contact and left unperturbed for 10 minutes. Afterwards, both cells were aspirated by pipettes, and the negative pressure in one of the pipettes (holding pipette) was adjusted to hold one cell firmly. The pressure in the other pipette was then increased in a stepwise fashion and at each step a separation attempt was performed, which involved moving the pulling pipette away from the holding pipette with a constant speed of 20 μm/s up to a distance of 20 μm. Pressure was recorded at each separation attempt, and subsequently separation force (*F_s_*) was calculated according to the equation:

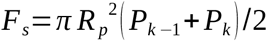

where *R_p_* is the pulling pipette radius, *k* =1,2,… is the attempt number and *P*_*k*−1_ and *P_k_* are pressure values in the last unsuccessful and the first successful separation attempt, respectively. Experiments where more than 6 attempts were needed for separation were excluded from the study to avoid mechanosensitive stiffening of the separated cells.

### Transmission Electron Microscopy (TEM)

For high-pressure freezing of cells, sapphire disks of 1.4 x 0.05 mm in diameter (Wohlwend, Sennwald, CH) were carbon-coated to a thickness of 10 nm using the Leica EM ACE600 high-vacuum coating device (Leica Microsystems). The pattern of a Maxtaform H15 finder grid (Science Services, LF 135-Ni) was evaporated onto the disk surface, and the coat was stabilized by baking overnight at 120°C. After plasma-cleaning for 2 min (Harrick plasma cleaner, RF level Medium), sapphire disks were incubated overnight at 4°C in 10 μM solution of Concavalin A (Sigma-Aldrich) and washed thoroughly in PBS. They were then placed into cup-shaped aluminium planchettes with cavity dimensions of 2 mm inner diameter and 100 μm indentation (Wohlwend, Sennwald, CH). Primary progenitor cells were prepared as described in ‘Cell culture’ above and plated onto the sapphires with cells from 1 embryo in average distributed over 2 disks. Cells were allowed to form spontaneous contacts and adhere to the disk surface for 10 min at RT. 1 μl of 5% BSA (Sigma-Aldrich, A-9647) in medium equilibrated to RT was then added as space filler and anti-freezing agent. The flat side of an aluminium planchette with a 300 μm indentation was used as lid and excess of solution was removed with filter paper. The sandwiched samples were high-pressure frozen instantaneously using the using the HPM-010 high pressure freezing machine (Leica Microsystems), transferred to cryo-vials (Biozym, T311-2) and then stored in liquid nitrogen.

For freeze-substitution, samples were processed in an AFS1 device (Leica Microsystems) with ethanol in the loading chamber. Two substitution cocktails were applied consecutively: (i) 1% tannic acid (Sigma-Aldrich, 403040) in non-hydrous acetone (VWR, 8.22251) and (ii) 1% osmium (EMS, 19134) plus 0.2% uranyl acetate (20% stock in methanol; AL-Labortechnik, 77870.2) in non-hydrous acetone. 2 ml screw-cap Nalgene® cryo-vials (Sigma-Aldrich, V4632) were used for substitution, filled with 1 ml of cocktail. The sequence for infiltration and stepwise warming was as follows: 24 h incubation in 0.1% tannic acid in acetone at −82°C, 3 times 10 min wash in acetone at −82°C, 6 h incubation in 1% osmium plus 0.2% uranyl acetate in acetone at −82°C, temperature rise 15°C/h to −60°C, 3 h incubation at −60°C, temperature rise 15°C/h to −30°C, 3 h incubation at −30°C, 3 times 10 min wash in acetone at −30°C, temperature rise 15°C/h to 4°C. Sapphires were then removed from the aluminium planchettes and embedded in epoxy resin (Durcupan® ACM, Fluca). Samples were consecutively infiltrated with a 3:1 mixture of acetone and Durcupan® for 1 h at 4°C, 1:1 acetone/Durcupan® for 1.5 h at 4°C, 1:3 aceton/Durcupan® for 2 h at 4°C and mere Durcupan® overnight at RT. Samples were transferred to BEEM capsules (EMS, 70020-B) filled with freshly prepared Durcupan® and cured for 48 h at 60°C in an oven. Serial ultrathin sections (70-80 nm) were cut using an UC7 ultramicrotome (Leica Microsystems) and collected onto formvar-coated copper slot grids. The sections were then contrast enhanced by incubating them in 1% aqueous uranyl acetate for 10 min at RT and Reynold’s lead citrate for 2 min at RT.

Sections were examined in a Tecnai 10 transmission electron microscope (Thermo Fisher Scientific) operated at 80 kV and equipped with an EMSIS side mounted camera Megaview III (Münster, Germany). Images were processed with Radius software (EMSIS) and Photoshop (Adobe) without changing any specific feature. For high-resolution analysis, sections were examined in a JEM 2800 scanning-TEM (Jeol) operated at 200 kV in STEM bright-field mode, and equipped with a side-mounted OSIS Veleta camera (EMSIS).

### Correlative Light and Electron Microscopy (CLEM)

Disks of 1 cm in diameter were cut from Aclar® foil (thickness 198 μm; TedPella, 10501-10) and placed in sterile Corning ® 12-well plates (Sigma-Aldrich, CLS 3737). Dissociated cells were plated on the these disks (cells from 1 blastoderm /disk) and allowed to form spontaneous cell-cell contacts and adhere to the disk surface for 10 min at RT. Cells were then fixed with 4% paraformaldehyde (PFA, Sigma-Aldrich, 158127) in PBS (pH 7.4) for 10 min at RT and washed 3x with PBS. Subsequently, Phalloidin conjugated with Alexa488 (Invitrogen, 1/250 in PBT) was applied to the cells for 3h at RT to label F-Actin. In an additional round of fixation, 4% PFA plus 0.05% glutaraldehyde (Agar Scientific, R1020) in PBS was applied for 20 min at RT. After washing in PBS, 50 mM glycine (VWR, 24403.298) in PBS was used to block free aldehyde groups for 20 min at RT. After washing in PBS again, samples were dehydrated in graded ethanol (50%, 70%, 90%, 96%, 100%) and embedded in LR-White resin (Sigma, 62661). Samples were consecutively infiltrated with a 1:1 mixture of ethanol to LR-White, 1:2 ethanol/LR-White and mere LR-White for 20 min each at 4°C. Samples were transferred to gelatin capsules (Science Services, 70103), filled with fresh LR-White, capped tightly and cured for 12 h at 50°C in an oven. Sections were cut at 180 nm and mounted on 15 mm glass coverslips coated with a Tissue Capture Pen (EMS, 71314-10). Sections were then embedded in VectaShield (Vector Laboratories), coverslipped and imaged under a LSM 880 microscope (Zeiss) with an oil immersion objective (40x NA 1.4) using an ‘Airy Scan’ detector. Overview images were taken to facilitate localization of doublets on the section. After fluorescence imaging, coverslips were removed from glass slides, and sections contrast enhanced by incubating them in 1% aqueous uranyl acetate for 10 min at RT and Reynold’s lead citrate for 4 min at RT. Sections were then observed under a Merlin VP Compact FE-Scanning Electron Microscope (Zeiss) using an In-lens Duo detector (In-lens SE and In-lens BSE). Images from details were first aligned to the overviews where the whole cell-cell doublet was visible using SIFT algorithm^55^. Subsequently, fluorescent images were aligned with the EM overviews using eC-CLEM method^56^ implemented in Icy open source software^58^.

### Atomic Force Microscopy (AFM)

Cell-cortex tension measurements on single cells were performed as described previously^29^, with preparation of single cells as described in ‘Cell culture’ section. For each experiment, individual cells from five blastoderm preparations were seeded on a tissue culture dish with cover glass bottom (FluoroDish) containing DMEM/F12 either alone (control) or complemented with 5 nM, 50 nM LPA or 10 μM Bb. Cells were probed using an AFM (NanoWizard 4^®^ BioScience, JPK Instruments) mounted on an inverted fluorescent microscope (Olympus IX71). Commercial colloidal force probes (CP-qp-CONT-BSA-A, NanoAndMore USA) were passivated with heat-inactivated fetal calf serum (FCS, Invitrogen) for 1h at room temperature to avoid non-specific adhesion of the bead to the cells. Force-distance curves were acquired using 500 pN contact force and 1 μm s^−1^ approach/retract velocity. Up to three curves with 10 s waiting time between successive curves were taken per cell to prevent any history effect. Indentation was calculated from the tip displacement. To obtain the values of cell-cortex tension, the liquid droplet model was applied as described previously^29^, with the following adjustments: for determining cell-cortex tension, a force versus indentation line-fit between 200 nm and 300 nm indentation range was applied.

### Reagents and inhibitors

Fetal BSA (GIBCO), heat-passivated FBS (Fetal Bovine Serum, Invitrogen), heat-inactivated FCS (Fetal Calf Serum, Invitrogen), and 1-Oleoyl lysophosphatidic acid (LPA, Tocris Bioscience) were used at the indicated concentrations (1 nM, 5 nM, 10 nM and 50 nM, respectively). Pharmacological inhibitors were used at the following concentrations: 10 μM active (−) or inactive Blebbistatin (+) (Tocris Bioscience), 0.3 μM Latrunculin-B (Sigma-Aldrich), 100 nM Jasplakinolide (Invitrogen), and 2 mg/mL Concavalin A (Sigma-Aldrich).

### Polymer microwell preparation

To facilitate imaging of the cell-cell contacts in the focal plane, a microwell setup was used as described^16^. In order to ascertain that the cell doublet will always remain in the correct position during the experiment, microwells with a range of well diameters (15-30 μm) and 50 μm depth were prepared. PDMS stamps containing the negative of the desired pattern were gently pressed to droplets of My Polymer 134 (My Polymers) applied to Mattek glass bottom petri dishes and thenUV-curated (Thorlabs UV LED 365nm) in nitrogen atmosphere for up to 1h, at which point the PDMS stamps were peeled off.

### Statistical analysis and repeatability of experiments

Statistical analysis of data was performed using the GraphPad Prism 6 software and statsmodel python package. Statistical details of experiments are reported in the figures and figure legends. To test for normality of a sample, a D’Agostino & Pearson omnibus normality test was used. In case two samples were compared and normal distribution was assumed, an unpaired *t*-test was performed, while Mann-Whitney test was performed in case of not normally distributed data. In case more than two normally distributed samples were compared, an ANOVA was performed followed by Tukey’s multiple comparison test. Alternatively, *t-*test was performed with Bonferroni correction for multiple comparisons as stated in detail in the figure legends. If no normal distribution could be assumed, a Kruskal-Wallis test followed by Dunn’s multiple comparison test was used. At least more than three independent experiments (N) were performed unless stated otherwise in the figure legend. No statistical method was used to predetermine sample size, the experiments were not randomized and the investigators were not blinded to allocation during experiments and outcome assessment. *P*-value of < 0.05 was considered as significant.

## Acknowledgements

We would like to thank Edouard Hannezo for discussions, Shayan Shami Pour and Daniel Capek for help with data analysis, Vanessa Barone and other members of the Heisenberg laboratory for thoughtful discussions and comments on the manuscript. We also thank Jack Merrin for preparing the microwells, and the Scientific Service Units at IST Austria, specifically Bioimaging and Electron Microscopy, and the Zebrafish Facility for continuous support. We acknowledge Hitoshi Morita for the kind gift of VinculinB-GFP plasmid. This research was supported by an ERC Advanced Grant (MECSPEC) to C.-P.H, EMBO Long Term grant (ALTF 187-2013) to M.S and IST Fellow Marie-Curie COFUND No. P_IST_EU01 to J.S.

## Author information

## Contributions

J.S., M.S., and C.-P.H. developed and designed the research. J.S. and M.S. contributed equally to this project and performed most of the experiments. S.C.-M. performed and analyzed the AFM experiments., W.A.K. performed CLEM experiments and ultra-structural analysis, K.H. and S.F.G.K. helped analyse results and participated in study design. J.S., M.S., and C.-P.H. wrote the manuscript. C.-P.H supervised the work.

## Additional information

Supplementary information is available in the online version of the paper.

## Supplementary Information

**Figure S1.**
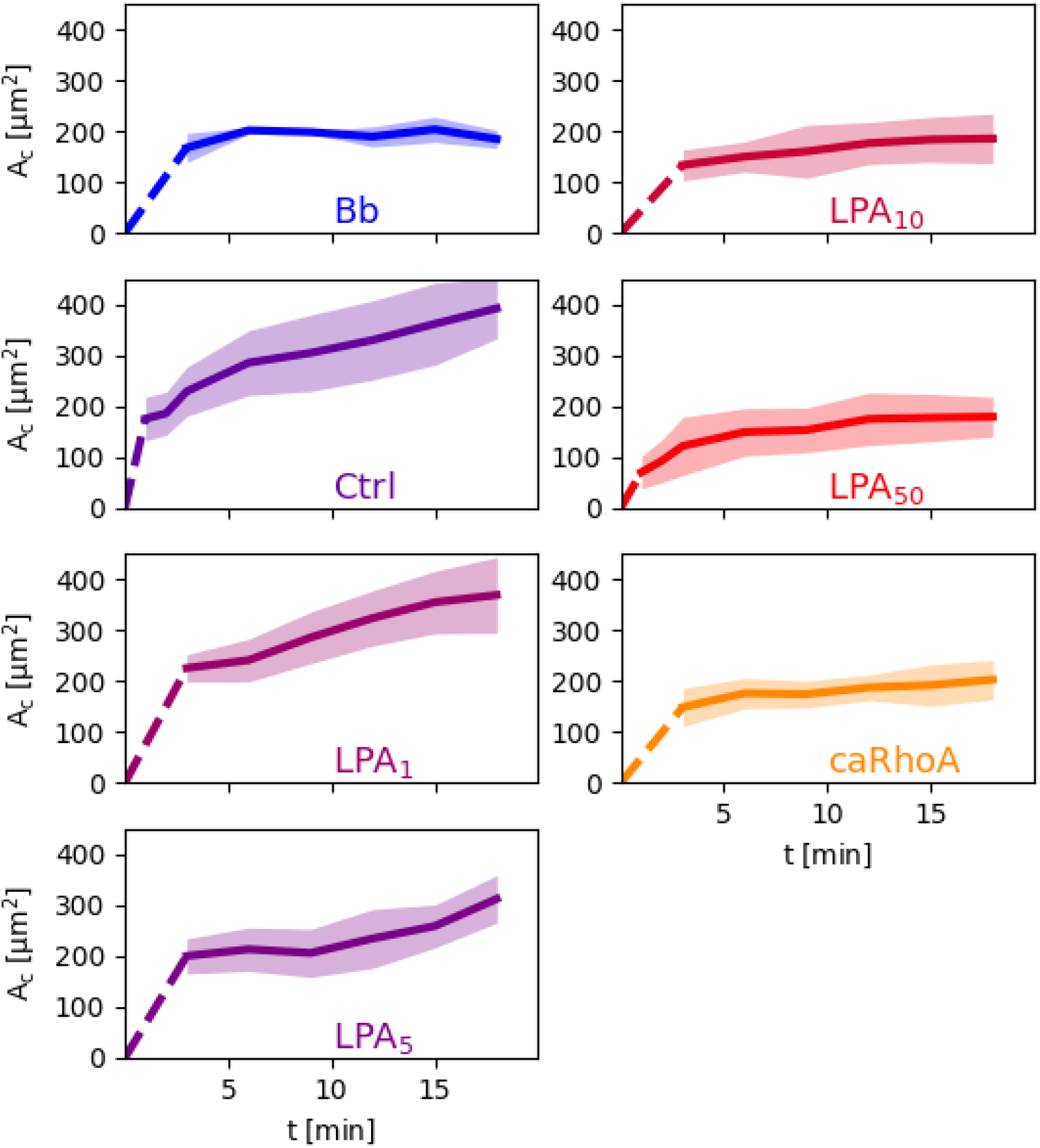
Cell-cell contact size as a function of time in control doublets and doublets exposed to 10 μM Bb or various concentrations of LPA (1-50 nM), or overexpressing caRhoA. Shadowed areas denote standard deviation. Dashed lines connect contact formation (0 min) with the first time point when data were collected. N and n for Ctrl, Bb, LPA_5_ and LPA_50_ as defined in Fig. 1. (LPA_1_) N = 3, n = (3min: 7, 6min: 7, 9min: 7, 12min: 7, 15min: 7, 18min: 7); (caRhoA) N = 3, n = (3min: 18, 6min: 18, 9min: 18, 12min: 18, 15min: 18, 18min: 18); (LPA_10_) N = 4, n = (3min: 12, 6min: 12, 9min: 9, 12min: 12, 15min: 8, 18min: 9)

**Figure S2.**
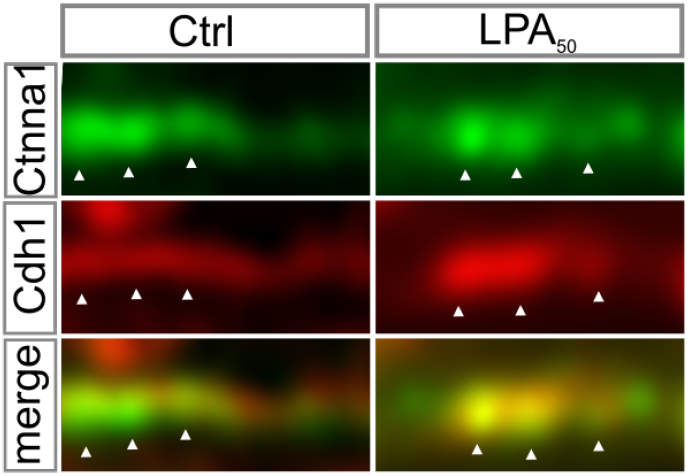
Cadherin-Catenin co-localization at the contact edge of control (left column) and LPA (50 nM)-treated (right column) progenitor cell doublets. High-resolution (‘Airy Scan’) images showing endogenous Ctnna1 (upper row, green), E-cadherin/Cdh1 (middle row, red) and merged (lower row) localization. Images were taken 30 min after contact formation. White arrowheads point at individual Cadherin-Catenin clusters.

**Figure S3.**
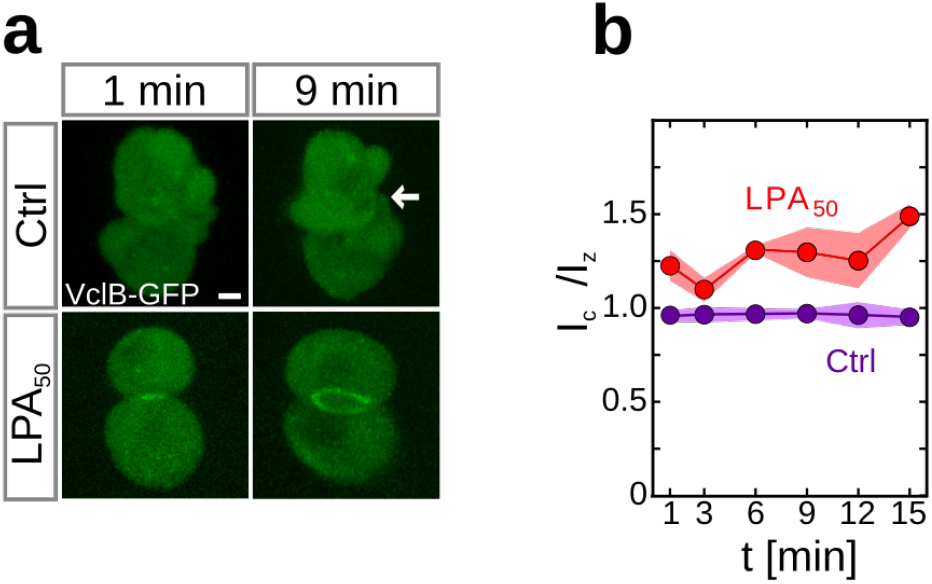
(a) Fluorescence images of cell doublets expressing GFP-tagged VinculinB in the presence or absence of LPA_50_. Arrow points at cell-cell contact lacking VinculinB-GFP accumulation. **(b)** Rim to centre intensity ratios of VinculinB-GFP in control (pink) and LPA_50_-treated (red) doublets. Shadowed areas denote standard deviations. (Ctrl) N = 3, n = (1min: 24, 3min: 20, 6min: 20, 9min: 20, 12min: 20, 15min: 20); (LPA_50_) N = 3, n = (1min: 4, 3min:4, 6min: 4, 9min: 4, 12min: 3, 15min: 3)

**Figure S4.**
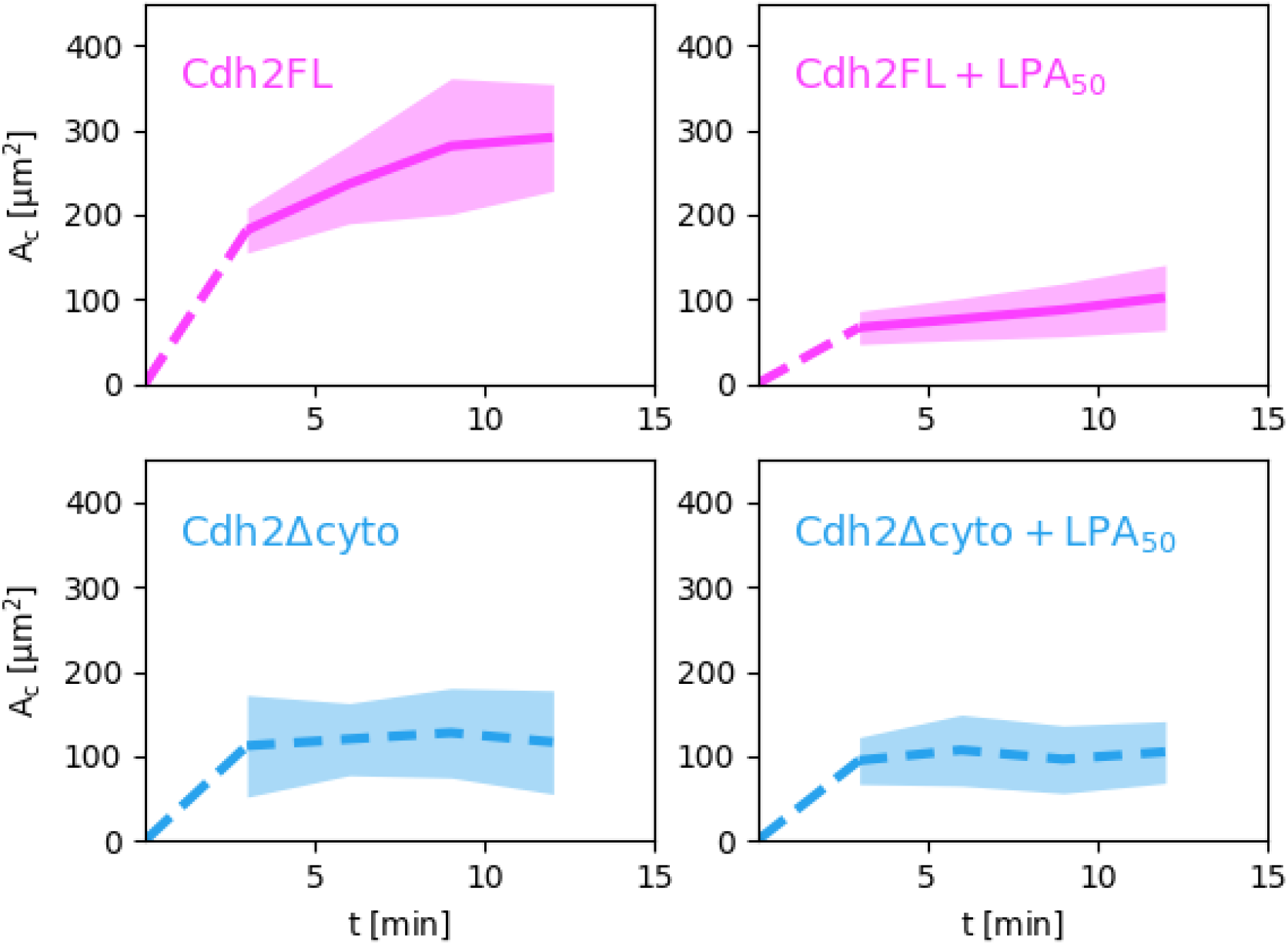
Cell-cell contact size as a function of time in progenitor cell doublets overexpressing full-length Cdh2 (Cdh2FL, pink) or Cdh2 lacking its cytoplasmic C-terminus (Cdh2 *Δ* cyto, blue) in the presence (dashed line) or absence (solid line) of 50 nM LPA. Shadowed areas denote standard deviation. Dashed lines connect contact formation (0 min) with the first time point when data were collected. N and n as stated in Fig. 5 c.

## References

1. Maitre, J.-L. et al. Adhesion Functions in Cell Sorting by Mechanically Coupling the Cortices of Adhering Cells. Science 338, 253–256 (2012).

2. Winklbauer, R. Cell adhesion strength from cortical tension - an integration of concepts. J. Cell Sci. 128, 3687–3693 (2015).

3. Yoshida, C. & Takeichi, M. Teratocarcinoma cell adhesion: identification of a cell-surface protein involved in calcium-dependent cell aggregation. Cell 28, 217–224 (1982).

4. Hong, S., Troyanovsky, R. B. & Troyanovsky, S. M. Cadherin exits the junction by switching its adhesive bond. J. Cell Biol. 192, 1073–1083 (2011).

5. Hinck, L. Dynamics of cadherin/catenin complex formation: novel protein interactions and pathways of complex assembly. J. Cell Biol. 125, 1327–1340 (1994).

6. Rimm, D. L., Koslov, E. R., Kebriaei, P., Cianci, C. D. & Morrow, J. S. Alpha 1(E)-catenin is an actin-binding and -bundling protein mediating the attachment of F-actin to the membrane adhesion complex. Proc. Natl. Acad. Sci. U. S. A. 92, 8813–8817 (1995).

7. Nagafuchi, A. The roles of catenins in the cadherin-mediated cell adhesion: functional analysis of E-cadherin-alpha catenin fusion molecules. J. Cell Biol. 127, 235–245 (1994).

8. Maître, J.-L. & Heisenberg, C.-P. Three functions of cadherins in cell adhesion. Curr. Biol. 23, R626–33 (2013).

9. Steinberg, M. S. Differential adhesion in morphogenesis: a modern view. Curr. Opin. Genet. Dev. 17, 281–286 (2007).

10. Stachowiak, J. C. et al. Membrane bending by protein-protein crowding. Nat. Cell Biol. 14, 944–949 (2012).

11. Perez, T. D., Tamada, M., Sheetz, M. P. & James Nelson, W. Immediate-Early Signaling Induced by E-cadherin Engagement and Adhesion. J. Biol. Chem. 283, 5014–5022 (2007).

12. Yamada, S. & James Nelson, W. Localized zones of Rho and Rac activities drive initiation and expansion of epithelial cell–cell adhesion. J. Cell Biol. 178, 517–527 (2007).

13. Anastasiadis, P. Z. et al. Inhibition of RhoA by p120 catenin. Nat. Cell Biol. 2, 637–644 (2000).

14. Borghi, N. et al. E-cadherin is under constitutive actomyosin-generated tension that is increased at cell-cell contacts upon externally applied stretch. Proc. Natl. Acad. Sci. U. S. A. 109, 12568–12573 (2012).

15. Roh-Johnson, M. et al. Triggering a cell shape change by exploiting preexisting actomyosin contractions. Science 335, 1232–1235 (2012).

16. Engl, W., Arasi, B., Yap, L. L., Thiery, J. P. & Viasnoff, V. Actin dynamics modulate mechanosensitive immobilization of E-cadherin at adherens junctions. Nat. Cell Biol. 16, 587–594 (2014).

17. Buckley, C. D. et al. Cell adhesion. The minimal cadherin-catenin complex binds to actin filaments under force. Science 346, 1254211 (2014).

18. Sim, J. Y. et al. Spatial distribution of cell–cell and cell–ECM adhesions regulates force balance while maintaining E-cadherin molecular tension in cell pairs. Molecular Biology of the Cell 26, 2456–2465 (2015).

19. Leerberg, J. M. et al. Tension-sensitive actin assembly supports contractility at the epithelial zonula adherens. Curr. Biol. 24, 1689–1699 (2014).

20. Thomas, W. A. et al. α-Catenin and vinculin cooperate to promote high E-cadherin-based adhesion strength. J. Biol. Chem. 288, 4957–4969 (2013).

21. Yonemura, S., Wada, Y., Watanabe, T., Nagafuchi, A. & Shibata, M. alpha-Catenin as a tension transducer that induces adherens junction development. Nat. Cell Biol. 12, 533–542 (2010).

22. Desai, R. et al. Monomeric α-catenin links cadherin to the actin cytoskeleton. Nat. Cell Biol. 15, 261–273 (2013).

23. Hirano, S., Kimoto, N., Shimoyama, Y., Hirohashi, S. & Takeichi, M. Identification of a neural alpha-catenin as a key regulator of cadherin function and multicellular organization. Cell 70, 293–301 (1992).

24. Yao, M. et al. Force-dependent conformational switch of α-catenin controls vinculin binding. Nat. Commun. 5, 4525 (2014).

25. Huveneers, S. et al. Vinculin associates with endothelial VE-cadherin junctions to control force-dependent remodeling. J. Cell Biol. 196, 641–652 (2012).

26. le Duc, Q. et al. Vinculin potentiates E-cadherin mechanosensing and is recruited to actin-anchored sites within adherens junctions in a myosin II-dependent manner. J. Cell Biol. 189, 1107–1115 (2010).

27. Mills, G. B. & Moolenaar, W. H. The emerging role of lysophosphatidic acid in cancer. Nat. Rev. Cancer 3, 582–591 (2003).

28. Takesono, A. et al. Solute carrier family 3 member 2 (Slc3a2) controls yolk syncytial layer (YSL) formation by regulating microtubule networks in the zebrafish embryo. Proc. Natl. Acad. Sci. U. S. A. 109, 3371–3376 (2012).

29. Krieg, M. et al. Tensile forces govern germ-layer organization in zebrafish. Nat. Cell Biol. 10, 429–436 (2008).

30. Puech, P.-H., Poole, K., Knebel, D. & Muller, D. J. A new technical approach to quantify cell-cell adhesion forces by AFM. Ultramicroscopy 106, 637–644 (2006).

31. Panther, E. et al. The influence of lysophosphatidic acid on the functions of human *dendritic cells*. J. Immunol. 169, 4129–4135 (2002).

32. Cavey, M., Rauzi, M., Lenne, P.-F. & Lecuit, T. A two-tiered mechanism for stabilization and immobilization of E-cadherin. Nature 453, 751–756 (2008).

33. Truong Quang, B.-A. et al. Principles of E-Cadherin Supramolecular Organization In Vivo. Curr. Biol. 23, 2197–2207 (2013).

34. Kale, G. R. et al. Distinct contributions of tensile and shear stress on E-cadherin levels during morphogenesis. Nat. Commun. 9, 5021 (2018).

35. Dufour, S., Mège, R.-M. & Thiery, J. P. α-catenin, vinculin, and F-actin in strengthening E-cadherin cell-cell adhesions and mechanosensing. Cell Adh. Migr. 7, 345–350 (2013).

36. Bianchini, J. M. et al. Reevaluating αE-catenin monomer and homodimer functions by characterizing E-cadherin/αE-catenin chimeras. J. Cell Biol. 210, 1065–1074 (2015).

37. Salbreux, G., Charras, G. & Paluch, E. Actin cortex mechanics and cellular morphogenesis. Trends in Cell Biology 22, 536–545 (2012).

38. Bertocchi, C. et al. Nanoscale architecture of cadherin-based cell adhesions. Nat. Cell Biol. 19, 28–37 (2017).

39. Hong, S., Troyanovsky, R. B. & Troyanovsky, S. M. Binding to F-actin guides cadherin cluster assembly, stability, and movement. J. Cell Biol. 201, 131–143 (2013).

40. Bertet, C., Sulak, L. & Lecuit, T. Myosin-dependent junction remodelling controls planar cell intercalation and axis elongation. Nature 429, 667–671 (2004).

41. Collins, C. & James Nelson, W. Running with neighbors: coordinating cell migration and cell–cell adhesion. Curr. Opin. Cell Biol. 36, 62–70 (2015).

42. Maruthamuthu, V. & Gardel, M. L. Protrusive activity guides changes in cell-cell tension during epithelial cell scattering. Biophys. J. 107, 555–563 (2014).

43. Hoffman, B. D. & Yap, A. S. Towards a Dynamic Understanding of Cadherin-Based Mechanobiology. Trends Cell Biol. 25, 803–814 (2015).

44. Leckband, D. E. & de Rooij, J. Cadherin adhesion and mechanotransduction. Annu. Rev. Cell Dev. Biol. 30, 291–315 (2014).

45. Yap, A. S., Duszyc, K. & Viasnoff, V. Mechanosensing and Mechanotransduction at Cell-Cell Junctions. Cold Spring Harb. Perspect. Biol. 10, (2018).

46. Pinheiro, D. & Bellaϊche, Y. Mechanical Force-Driven Adherens Junction Remodeling and Epithelial Dynamics. Dev. Cell 47, 391 (2018).

47. Westerfield, M. The Zebrafish Book: A Guide for the Laboratory Use of Zebrafish (Danio Rerio). (2007).

48. Kimmel, C. B., Ballard, W. W., Kimmel, S. R., Ullmann, B. & Schilling, T. F. Stages of embryonic development of the zebrafish. Dev. Dyn. 203, 253–310 (1995).

49. Choe, C. P. et al. Wnt-dependent epithelial transitions drive pharyngeal pouch formation. Dev. Cell 24, 296–309 (2013).

50. Behrndt, M. et al. Forces driving epithelial spreading in zebrafish gastrulation. Science 338, 257–260 (2012).

51. Chu, Y.-S. et al. Force measurements in E-cadherin-mediated cell doublets reveal rapid adhesion strengthened by actin cytoskeleton remodeling through Rac and Cdc42. J. Cell Biol. 167, 1183–1194 (2004).

52. Arboleda-Estudillo, Y. et al. Movement directionality in collective migration of germ layer progenitors. Curr. Biol. 20, 161–169 (2010).

53. Rueden, C. T. et al. ImageJ2: ImageJ for the next generation of scientific image data. BMC Bioinformatics 18, 529 (2017).

54. Banovic, D. et al. Drosophila neuroligin 1 promotes growth and postsynaptic differentiation at glutamatergic neuromuscular junctions. Neuron 66, 724–738 (2010).

55. Lowe, D. G. Distinctive Image Features from Scale-Invariant Keypoints. International Journal of Computer Vision 60, 91–110 (2004).

56. Heiligenstein, X., Paul-Gilloteaux, P., Belle, M., Raposo, G. & Salamero, J. eC-CLEM: Flexible Multidimensional Registration Software for Correlative Microscopies with Refined Accuracy Mapping. Microscopy and Microanalysis 23, 360–361 (2017).

57. de Chaumont, F. et al. Icy: an open bioimage informatics platform for extended reproducible research. Nat. Methods 9, 690–696 (2012).

